# Occurrence of *Amaranthus palmeri* in Israeli agriculture: status of spread and response to glyphosate and trifloxysulfuron

**DOI:** 10.1101/2023.07.25.550449

**Authors:** Jackline Abu-Nassar, Amit Wallach, Eilon Winkler, Hanan Eizenberg, Maor Matzrafi

## Abstract

*Amaranthus palmeri* S. Watson is a dioecious annual weed species, originating from the Southern USA, spreading rapidly beyond its original range into Europe and the Mediterranean region. In Israel, *A. palmeri* distribution has expanded quickly in recent years, with farmers reporting on weed control failure using acetolactate synthase (ALS) inhibitors. Furthermore, recent studies have documented glyphosate-resistant cases from other countries in the region, such as Spain, Greece, and Turkey. We conducted a survey in order to understand *A. palmeri* distribution and study the occurrence of herbicide resistance to both glyphosate and trifloxysulfuron in different fields across the country. According to our data, *A. palmeri* population locations are aligned with the major agricultural areas for summer field crops, including the Hula Valley, Jezreel Valley, and the Southern Coastal Plain. Regarding herbicide responses, while several populations showed a reduced response to glyphosate, dose-response assays did not show resistance to the recommended labeled field rate. For the ALS inhibitor trifloxysulfuron, the proportion of resistant individuals was very high, especially in the southern coastline region, with an R-value of 0.77. Four populations used for dose-response studies were highly resistant, surviving at four times the recommended labeled field rate of trifloxysulfuron (30 g a.i. ha^-1^). Sequencing of the ALS gene, Trp574 to Leu alteration in resistant populations was recorded. The high level of resistance observed in this study, alongside the target-site mutation found in populations of *A. palmeri*, endangers the future use of ALS inhibitors in corn, cotton, and other summer crops grown in Israel.

## 1 INTRODUCTION

In recent decades, we have seen increased reports of invasive weed species due to significant human-induced global change. Major drivers of this trend are import–export trades and the secondary climate change (Hübner et al., 2022; Matzrafi et al., 2021). *Amaranthus palmeri* S. Watson is a dioecious annual, erect, broadleaved weed species native to the southern USA, spreading rapidly into North America, Europe, and the Mediterranean region (Rubin & Matzrafi, 2015). *Amaranthus palmeri* is highly competitive against major summer field crops; even at low densities, its presence in the field may result in severe yield losses. According to a 2017 and 2019 national survey by the Weed Science Society of America (Van Wychen, 2017), *A. palmeri* is one of the most devastating weeds in maize and sorghum fields in the US. Since this species is a prolific seed producer, the presence of only a few female plants in the field may lead to substantial infestations during subsequent growing seasons (Norsworthy et al., 2014). Due to its high genetic variation, *A. palmeri* populations are particularly responsive to selection pressure (Yanniccari et al., 2022). This species is among the most infamous weed species for developing herbicide resistance patterns against multiple modes of action (MOAs) (Heap, 2023).

Currently, several *A. palmeri* populations have been identified as herbicide-resistant worldwide. Resistance has evolved to nine MOAs, including acetolactate synthase (ALS; Group 2), photosystem II (PSII; Group 5), protoporphyrinogen oxidase (PPO; Group 14), 4–hydroxyphenyl–pyruvate–dioxygenase (HPPD; Group 27), 5– enolpyruvyl shikimate–3–phosphate synthase (EPSPS; Group 9) inhibitors, and more (Heap, 2023). However, analyzing the variation among herbicide resistance reported cases, according to the different MOAs, shows that most of the reports originated outside of the Americas and are not related to EPSPS inhibitors. Only one report from South Africa identified an *A. palmeri* population showing resistance to both ALS and EPSPS inhibitors (Heap, 2023). In Europe, to date, most of the reported cases describe resistance to ALS inhibitors, with the exception of a report on glyphosate-resistant *A. palmeri* population from Spain (Manicardi *et al*., 2023). However, for both cases, it is assumed that herbicide resistance originated from the introduction of an herbicide-resistant population to the local flora and not as a response to selection pressure promoting the evolution of herbicide resistance in the reporting country.

In Israel, *A. palmeri* can be found in various habitats, such as field and vegetable crops, orchards, vineyards, roadsides, and disturbed sites (Rubin & Matzrafi, 2015). Herbarium samples were dated back to 1957, thus, this species was introduced to the Israeli flora over 60 years ago (Danin, 2022), when herbicide resistance to ALS and EPSPS inhibitors had not yet been reported (Heap, 2023). The common practice for weed control of *A. palmeri* for the Israeli farmer in field crops is using ALS inhibitors, such as foramsulfuron, imazamox, pyrithiobac-sodium, rimsulfuron, and trifloxysulfuron (Rubin & Matzrafi, 2015). Several PSII inhibitors, such as fluometuron, metribuzin, and prometryn are also used for weed management, but to a lesser extent.

In recent years, there has been an increase in reports of herbicide application failures to control *A. palmeri* in different habitats using ALS and EPSPS inhibitors. Although, in Israel, herbicide resistance was previously reported for ALS inhibitors (Heap, 2023), the extent of the phenomenon remains unknown. As ALS inhibitors are frequently used in various crops, it is valid to assume that herbicide resistance has evolved in multiple locations nationwide. Since glyphosate-resistant crops are not grown in Israel, glyphosate use is minimal but may be applied to control weeds between crop rows, avoiding the crop foliage using band spraying (Gerhards et al., 2022). Thus, the odds for glyphosate resistance evolution in field crops are very low, and, indeed, no glyphosate-resistant *A. palmeri* populations were previously documented in Israel. However, importing seeds as animal feed or for oil production from the US may have resulted in the introduction of glyphosate-resistant *A. palmeri* populations (Rubin & Matzrafi, 2015).

For this study, we collected seeds from 22 populations of *A. palmeri* from different locations across Israel. These seeds were germinated, and plants were tested for their response to varying rates of glyphosate and trifloxysulfuron. Resistance levels were further examined for a subset of populations showing reduced response to the herbicides in the initial test. The mechanism of resistance to the ALS inhibitor trifloxysulfuron was also studied.

## 2 MATERIALS AND METHODS

### 2.1 Distribution of *A. palmeri*

In order to better understand the distribution and occurrence of herbicide resistance throughout Israel, a detailed mapping of the infested sites, including a survey for the seed collection used for herbicide response studies, was performed during the spring-summer of 2020.

The mapping was performed using two methods: first, social media posts directed at local communities, farmers, agronomists, park rangers, and others who have sighted *A. palmeri*. These individuals were asked to access a link and report *A. palmeri* occurrences, including a specific location based on Google Maps, to a designated WhatsApp® group. After these resources reported a reasonable number of sites, we visited each location and confirmed *A. palmeri* identification. The second method involved scouting for *A. palmeri* in agricultural fields from the southern to the northern border of distribution. For each site, we recorded data on the cropping system and the specific crop grown in the field for the year of seed collection.

### 2.2 Seed collection method

*Amaranthus palmeri* seeds were collected using a sampling technique previously described by Culpepper et al. (2008) with minor adjustments. Seed collection includes randomly harvesting 4-6 seed heads from each plant for approximately 40 female plants spaced at least 10 m apart, collated haphazardly over the field. Across locations, the populations were separated by at least 5 km. Seeds from all females at each site were pooled and designated as a population. Seed heads were threshed and stored at 4°C until use.

### 2.3 Initial herbicide response screening

Seeds from all 22 populations were sown in trays filled with a commercial growth mixture (Tuf Marom-Golan®). Each tray contained 98 wells, with 5-8 seeds sown in each well. Upon emergence, seedlings were thinned to obtain one seedling per well. When most of the plants in each tray reached the 5-7 cm tall growth stage, trays were divided into three sections serving as untreated control, and plants that were sprayed using either half or the recommended labeled field rate. Herbicides used in this study were the EPSPS inhibitor - glyphosate (Roundup®, 36%, BASF) at the recommended labeled field rate of 720 g ae h^-1^ and the ALS inhibitor - trifloxysulfuron (Envok®, 75%, WG, Syngenta) at the recommended labeled field rate of 7.5 g ai h^-1^. Plants were sprayed using a chain-driven sprayer delivering 300 L ha^-1^ with a flat-fan 8001E nozzle (TeeJet®, Spraying Systems Co., Wheaton, IL, USA). Experiments were conducted at Newe-Ya’ar Research Center in a net-house during the natural season for *A. palmeri* growth at an average temperature of 33/22±3°C day/night and a photoperiod of 13-14 hours. Trays were placed on experimental benches and randomized every week. The survival rate was recorded 21 days after treatment (DAT). The maps used to describe survival rate were generated using ArcGIS Pro. 2.8.0. (ESRI, 2021).

### 2.4 Dose-response studies

The resistance level was investigated for a subset of populations that showed reduced herbicide efficacy, where plants had survived treatment with the recommended labeled field rate of a given herbicide. All the suspected herbicide-resistant populations were compared to a susceptible population chosen from the initial herbicide response assay. Plants were treated at 5-7 cm tall, with eight rates (0, 0.125X, 0.25X, 0.5X, 0.75X, 1X, 2X, and 4X), where X = recommended labeled field rate for each herbicide as specified above. All experiments were arranged in a completely randomized design, with 10 replicates for each treatment. Experiments were repeated twice. Shoot fresh weight and survival rate were recorded at 21 DAT.

### 2.5 Sequencing of the ALS gene

In order to detect structural substitutions, the ALS gene was sequenced and analyzed. Seeds from populations GENI, TZR, BDAR, YADB (ALS-resistant) and GE’A (ALS-sensitive), were sown in plastic trays filled with commercial growth mix. At the three-four leaf stage, trays were sprayed with 4X of trifloxysulfuron as described above. For each of the resistant populations, a leaf tissue was excised from each plant that have survived the treatment. DNA extraction was performed using the same procedures as described in Matzrafi *et al*. (2017). Primers that were used to sequence the ALS gene and the PCR procedure are described in Reinhardt *et al*. (2022). Sequence analyses and alignment were performed using the BioEdit software (Hall, 1999). The obtained sequences were compared to a known sequences of the ALS gene from *A. palmeri* (KY781923.1) obtained from NCBI (https://www.ncbi.nlm.nih.gov).

### 2.6 Statistical analyses

For the dose-response experiments, the effective dose that results in 50% inhibition of plant growth (GR_50_) and the lethal dose that resulted in 50% survival rate (LD_50_) were estimated for each population by fitting log-logistic three parameters curves to shoot fresh weight and survival rate as a function of herbicide dose (Seefeldt et al., 1995).

## 3 RESULTS

### 3.1 Distribution of *A. palmeri* populations within agricultural areas across Israel

*Amaranthus palmeri* populations were found in 22 locations within different agricultural habitats across Israel (Table 1). These locations are correlated with the major agricultural areas for summer field crops, including the Hula Valley, Jezreel Valley, and Southern Coastal Plain. The main crops in these regions are cotton, corn, and sorghum, as specified in Table 1. It is important to mention that these crops are grown as irrigated summer crops in Israel (Rubin & Matzrafi, 2015). In addition, two *A. palmeri* populations were recorded in olive and citrus orchards. Exploring the distribution of *A. palmeri*, we found that most plants were located within the crop rows. For two populations, plants were situated in Ag. adjacent habitats, GAD and KAV-4, and seeds were collected in the buffer zone within field margins. The agricultural habitat where plants were found only at field margins is a cotton field located at the Southern Coastal Plain for the TSH population (Table 1).

**Table 1.** Population ID and location across Israel. Description of the specific habitat and characteristics.

| **Pop ID** | **GPS coordinates** | **Habitat/cropping**<br>**system<sup>1</sup>** | **Specific**<br>**crop/habitat** | **Location of**<br>**plants<sup>2</sup>** |
| --- | --- | --- | --- | --- |
| 77R | 35.268918 ,32.770469 | Field Crops | Cotton | Within |
| AGAM | 35.596193 ,33.110525 | Field Crops | Peanut | Within |
| BAL | 35.440572 ,32.513936 | Orchards | Olive | Within |
| BDAR | 34.707144 ,31.826480 | Field Crops | Corn | Within |
| BHANA | 34.774417 ,31.936056 | Orchards | Citrus | Within |
| EVE | 35.105837 ,32.987606 | Field Crops | Corn | Within |
| GAD | 35.618915 ,33.027582 | Ag. adjacent | River Bank | Margins |
| GE'A | 34.590469 ,31.633119 | Field Crops | Watermelon | Within |
| GENI | 35.257735 ,32.660613 | Vegetable Crops | Tomato | Within |
| GON | 35.642584 ,33.122578 | Field Crops | Corn | Within |
| HAN | 35.252952 ,32.788345 | Field Crops | Sorghum | Within |
| KA'A | 35.192007 ,32.751109 | Vegetable Crops | Tomato | Within |
| KAV-4 | 35.616909 ,33.107714 | Ag. adjacent | Water channel | Margins |
| KMEN | 34.834727 ,31.746385 | Field Crops | Cotton | Within |
| OM | 34.982729 ,32.358332 | Field Crops | Peanut | Within |
| PAHA | 34.967676 ,32.488764 | Field Crops | Watermelon | Within |
| SIT | 34.958550 ,32.369460 | Field Crops | Peanut | Within |
| SVE | 34.938307 ,32.197163 | Vegetable Crops | Cabbage | Within |
| TSH | 34.894941 ,31.795957 | Field Crops | Cotton | Margins |
| TZR | 34.946995 ,31.762924 | Field Crops | Cotton | Within |
| YADB | 34.813900 ,31.801860 | Field Crops | Cotton | Within |
| YIZ | 35.345125 ,32.561502 | Field Crops | Sorghum | Within |

<sup>1</sup>Ag. Adjacent refers to agricultural habitats adjacent to natural habitats.
<sup>2</sup>The location where most of *A. palmeri* plants were located within the habitat, within
crop rows or in field margins.

### 3.2 Response of *A. palmeri* populations to glyphosate

As shown by the map in Fig. 1, populations can be clustered into four central regions across the country: the Galilee Panhandle, lower Galilee, and Coastal Plain-North and South. Although field crops may also be found in the southern Arava region, *A. palmeri* populations were not reported in this area (Matzrafi, personal observations). Plants from the 22 collected populations were tested for their response to two rates of glyphosate, half and full recommended labeled field rate. For half of the recommended field rate, several populations showed high survival rates, e.g., AGAM, EVE, GENI, and SIT (Fig. 1A). However, for the full rate, the percentage of survivors substantially decreased, with almost total control for all the tested populations (Fig. 1B).

**Figure 1.**
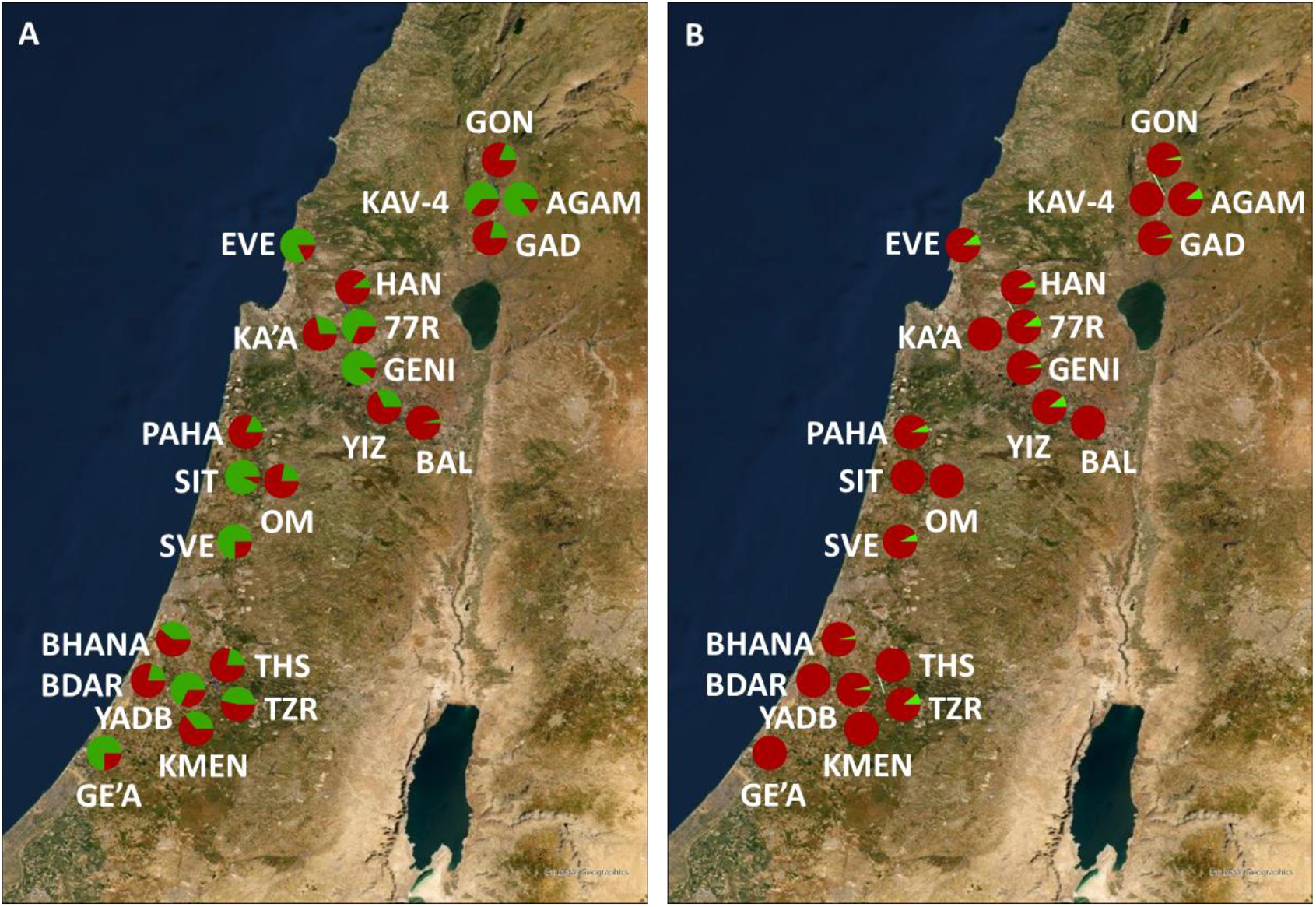
Map of *A. palmeri* populations, response to half (360 g ae ha^-1^; A) and the recommended labeled field rate (720 g ae ha^-1^; B) of glyphosate. In each population, the proportion of resistant (green) and susceptible (red) plants was based on the greenhouse screening of plants grown from field-collected seeds. n=28.

The resistance level was calculated for each population and at the regional scale (Table 2). For half of the recommended labeled field rate, the R proportion was high for plants from populations of AGAM (0.85), EVE (0.82), GENI (0.89), and SIT (0.93). The average R per region averaged 0.44 to 0.52. However, for the recommended labeled field rate, most of the populations were controlled in all regions. The highest proportion of resistant plants/populations was recorded in the lower Galilee region with an R per region value of 0.11 (Table 2).

**Table 2.** The proportion of plant survivors from 22 *A. palmeri* populations treated with half and the recommended labeled field rate of glyphosate.

| Pop<br>ID/Region | Half rate (360 g ae ha<sup>-1</sup>) |  |  |  | Full rate (720 g ae ha<sup>-1</sup>) |  |  |  |
| --- | --- | --- | --- | --- | --- | --- | --- | --- |
|  | No. | No. | R<sup>1</sup> | R per | No. | No. | R | R per |
|  | plants | plants |  | region | plants | plants |  | region |
|  | treated | surviving |  |  | treated | surviving |  |  |
| Galilee Panhandle |  |  |  |  |  |  |  |  |
| GON | 27 | 5 | 0.19 | 0.47 | 28 | 1 | 0.04 | 0.04 |
| GAD | 28 | 6 | 0.21 |  | 28 | 1 | 0.04 |  |
| KAV-4 | 28 | 18 | 0.64 |  | 28 | 0 | 0.00 |  |
| AGAM | 27 | 23 | 0.85 |  | 28 | 3 | 0.11 |  |
| Lower Galilee |  |  |  |  |  |  |  |  |
| EVE | 28 | 23 | 0.82 | 0.45 | 28 | 3 | 0.11 | 0.11 |
| YIZ | 28 | 9 | 0.32 |  | 28 | 3 | 0.11 |  |
| BAL | 28 | 1 | 0.04 |  | 21 | 0 | 0.00 |  |
| GENI | 28 | 25 | 0.89 |  | 28 | 1 | 0.04 |  |
| HAN | 28 | 3 | 0.11 |  | 27 | 2 | 0.07 |  |
| KA'A | 28 | 8 | 0.29 |  | 28 | 0 | 0.00 |  |
| 77R | 28 | 19 | 0.68 |  | 28 | 3 | 0.11 |  |
| Coastal plain-North |  |  |  |  |  |  |  |  |
| PAHA | 27 | 5 | 0.19 | 0.52 | 28 | 2 | 0.07 | 0.06 |
| OM | 27 | 6 | 0.22 |  | 28 | 0 | 0.00 |  |
| SIT | 28 | 26 | 0.93 |  | 28 | 0 | 0.00 |  |
| SVE | 28 | 21 | 0.75 |  | 27 | 2 | 0.07 |  |
| Coastal plain-South |  |  |  |  |  |  |  |  |
| TSH | 27 | 6 | 0.22 | 0.44 | 28 | 0 | 0.00 | 0.03 |
| BHANA | 28 | 11 | 0.39 |  | 28 | 1 | 0.04 |  |
| TZR | 28 | 13 | 0.46 |  | 28 | 3 | 0.11 |  |
| KMEN | 28 | 10 | 0.36 |  | 28 | 0 | 0.00 |  |
| BDAR | 28 | 6 | 0.21 |  | 27 | 0 | 0.00 |  |
| YADB | 28 | 19 | 0.68 |  | 27 | 1 | 0.04 |  |
| GE'A | 28 | 21 | 0.75 |  | 27 | 0 | 0.00 |  |

<sup>1</sup>R, population-level/region-level resistance to glyphosate is estimated as the proportion of survivors out of the total number of plants treated with the two herbicide rates.

### 3.3 Dose-response assays using glyphosate for a subset of *A. palmeri* populations

Further tests were carried out in order to validate the response of a subset of *A. palmeri* populations that were the least sensitive to glyphosate application. Populations EVE, YIZ, 77R, and TZR were tested for their response to increasing rates of glyphosate in comparison to population KAV-4, which is highly sensitive to the herbicide. Although the results of the two experimental runs were similar, they could not be combined due to statistical differences, and thus data are presented independently in Table S1.

For all tested populations, none of the plants survived the application of 1/2 the recommended labeled field rate of glyphosate (Fig S1A-E). Plants of the EVE population were found to be highly sensitive to glyphosate as no survivors were recorded even at 1/4 of the recommended labeled field rate, 180 g ae ha^-1^ (Fig S1G). Examination of both ED_50_ and LD_50_ showed very low values corresponding to most plants being highly affected using 1/4 of the recommended labeled field rate (Table S1). LD_50_ values were higher, showing that although plants were severely damaged, some survived the herbicide treatment. However, these values were still very low compared to the recommended labeled field rate used for glyphosate application in Israel.

### 3.4 Response of *A. palmeri* populations to trifloxysulfuron

In general, the efficacy for trifloxysulfuron was low and resistant populations were recorded in all four regions (Fig. 2). However, plants of populations originating from the southern Coastal Plain region had higher survival rates compared to the other regions for both half or the recommended labeled field rate of trifloxysulfuron.

**Figure 2.**
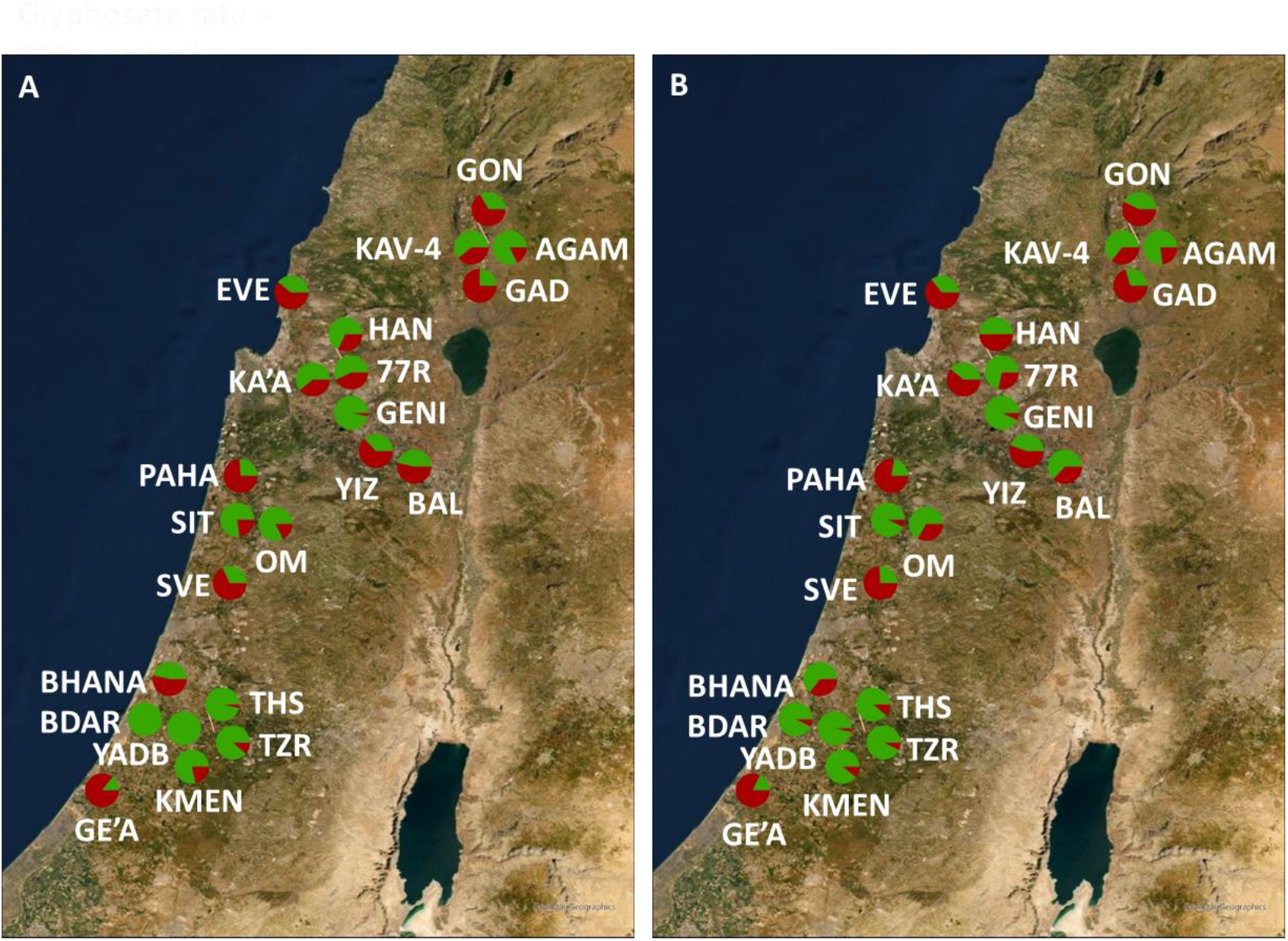
Map of *A. palmeri* populations, response to half (3.75 g ai ha^-1^; A) and the recommended labeled field rate (7.5 g ai ha^-1^; B) of trifloxysulfuron. In each population, the proportion of resistant (green) and susceptible (red) plants was based on the greenhouse screening of plants grown from field-collected seeds.

The proportion of resistant individuals was very similar among plants treated with half or the full recommended labeled field rate of trifloxysulfuron within each population (Fig. 2; Table 3). Also, the regional level of resistance was comparable for both rates, half and full, as shown by the R per region; Galilee Panhandle – 0.51 vs. 0.53, lower Galilee – 0.58 vs. 0.57, Coastal plain-North – 0.54 vs. 0.52, and Coastal plain-South 0.75 vs. 0.77. Populations tested for the recommended labeled field rate and scored a high R-value were mostly located in the Southern Coastal Plain region, TZR (0.93), BDAR (0.93), and YADB (0.96). Two other populations, one from each region (e.g., lower Galilee – GENI, northern Coastal plain – SIT), also showed high R per population with a value of 0.93 for plants treated with the recommended labeled field rate of trifloxysulfuron (Table 3).

**Table 3.** The proportion of plant survivors from 22 *A. palmeri* populations treated with half and the recommended labeled field rate of trifloxysulfuron.

|  | Half rate (3.75 g ai ha<sup>-1</sup>) |  |  |  | Full rate (7.5 g ai ha<sup>-1</sup>) |  |  |  |
| --- | --- | --- | --- | --- | --- | --- | --- | --- |
| Pop | No. | No. | R<sup>1</sup> | R per | No. | No. | R | R per |
| ID/Region | plants | plants |  | region | plants | plants |  | region |
|  | treated | surviving |  |  | treated | surviving |  |  |
| Galilee Panhandle |  |  |  |  |  |  |  |  |
| GON | 27 | 9 | 0.33 | 0.51 | 28 | 12 | 0.43 | 0.53 |
| GAD | 28 | 7 | 0.25 |  | 27 | 8 | 0.30 |  |
| KAV-4 | 28 | 17 | 0.61 |  | 28 | 18 | 0.64 |  |
| AGAM | 28 | 23 | 0.82 |  | 26 | 20 | 0.77 |  |
| Lower Galilee |  |  |  |  |  |  |  |  |
| EVE | 28 | 11 | 0.39 | 0.58 | 28 | 10 | 0.36 | 0.57 |
| YIZ | 27 | 10 | 0.37 |  | 27 | 12 | 0.44 |  |
| BAL | 28 | 13 | 0.46 |  | 27 | 17 | 0.63 |  |
| GENI | 28 | 27 | 0.96 |  | 28 | 26 | 0.93 |  |
| HAN | 28 | 19 | 0.68 |  | 28 | 14 | 0.50 |  |
| KA'A | 28 | 17 | 0.61 |  | 28 | 11 | 0.39 |  |
| 77R | 28 | 16 | 0.57 |  | 28 | 20 | 0.71 |  |
| Coastal plain-North |  |  |  |  |  |  |  |  |
| PAHA | 27 | 7 | 0.26 | 0.54 | 27 | 6 | 0.22 | 0.52 |
| OM | 28 | 23 | 0.82 |  | 27 | 18 | 0.67 |  |
| SIT | 26 | 20 | 0.77 |  | 28 | 26 | 0.93 |  |
| SVE | 28 | 9 | 0.32 |  | 27 | 7 | 0.26 |  |
| Coastal plain-South |  |  |  |  |  |  |  |  |
| TSH | 27 | 26 | 0.96 | 0.75 | 27 | 24 | 0.89 | 0.77 |
| BHANA | 28 | 13 | 0.46 |  | 28 | 18 | 0.64 |  |
| TZR | 26 | 23 | 0.88 |  | 28 | 26 | 0.93 |  |
| KMEN | 28 | 22 | 0.79 |  | 28 | 25 | 0.89 |  |
| BDAR | 28 | 28 | 1.00 |  | 28 | 26 | 0.93 |  |
| YADB | 28 | 28 | 1.00 |  | 28 | 27 | 0.96 |  |
| GE'A | 28 | 4 | 0.14 |  | 28 | 5 | 0.18 |  |

<sup>1</sup>R, population-level/region-level resistance to trifloxysulfuron estimated as the proportion of survivors out of the total number of plants treated with the two herbicide rates.

### 3.5 Dose-response assays using trifloxysulfuron for a subset of *A. palmeri* populations

Four populations suspicious as trifloxysulfuron-resistant were compared to a sensitive population using dose-response studies (Fig. 3). The experiment was conducted twice, however, due to statistical differences among experimental runs, data were not combined. A high level of resistance was recorded for all tested populations e.g. TZR, BDAR, YADB, and GENI (Fig 3, Fig S2). Since no correlation between shoot fresh weight and increasing trifloxysulfuron rate was observed, exponential decay models were not able to describe trifloxysulfuron response for any of the resistant populations (Fig S2). In addition, although a reduction in shoot fresh weight was observed, all treated plants survived tested rates of up to four times the recommended labeled field rate, 30 g ai ha^-1^.

**Figure 3.**
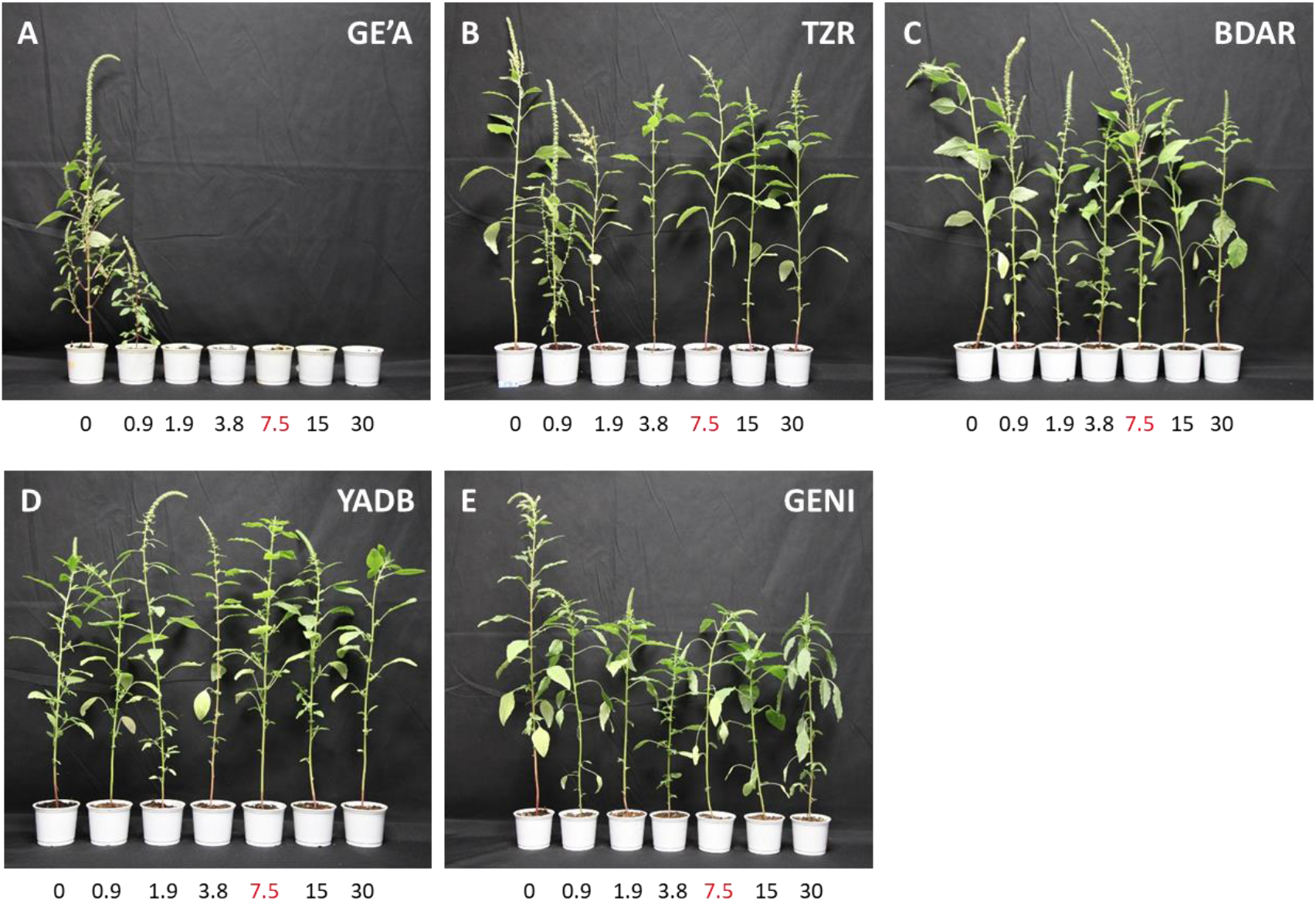
Representative images of trifloxysulfuron-treated *A. palmeri* plants from five populations. The recommended labeled field rate = 7.5 g ai ha^-1^, marked in red. n=10.

### 3.6 Mechanism of resistance to ALS inhibitors

The nucleotide sequence of the ALS gene was amplified using two pairs of primers (Reinhardt et al., 2022). Sequences were then aligned with each other and compared to the *A. thaliana* ALS gene, in order to identify the nucleotide substitutions leading to point mutations. For the initial part of the ALS gene, no substitutions were found in positions Pro197 and Ala205 (data not shown). A substitution in position 574 from Trp to Leu, was found in all populations (Fig. 4). Mutation frequency was higher for populations BDAR and YADB, however, further data is needed in order to undesratnd the frequency of this mutation in the population level.

**Figure 4.**
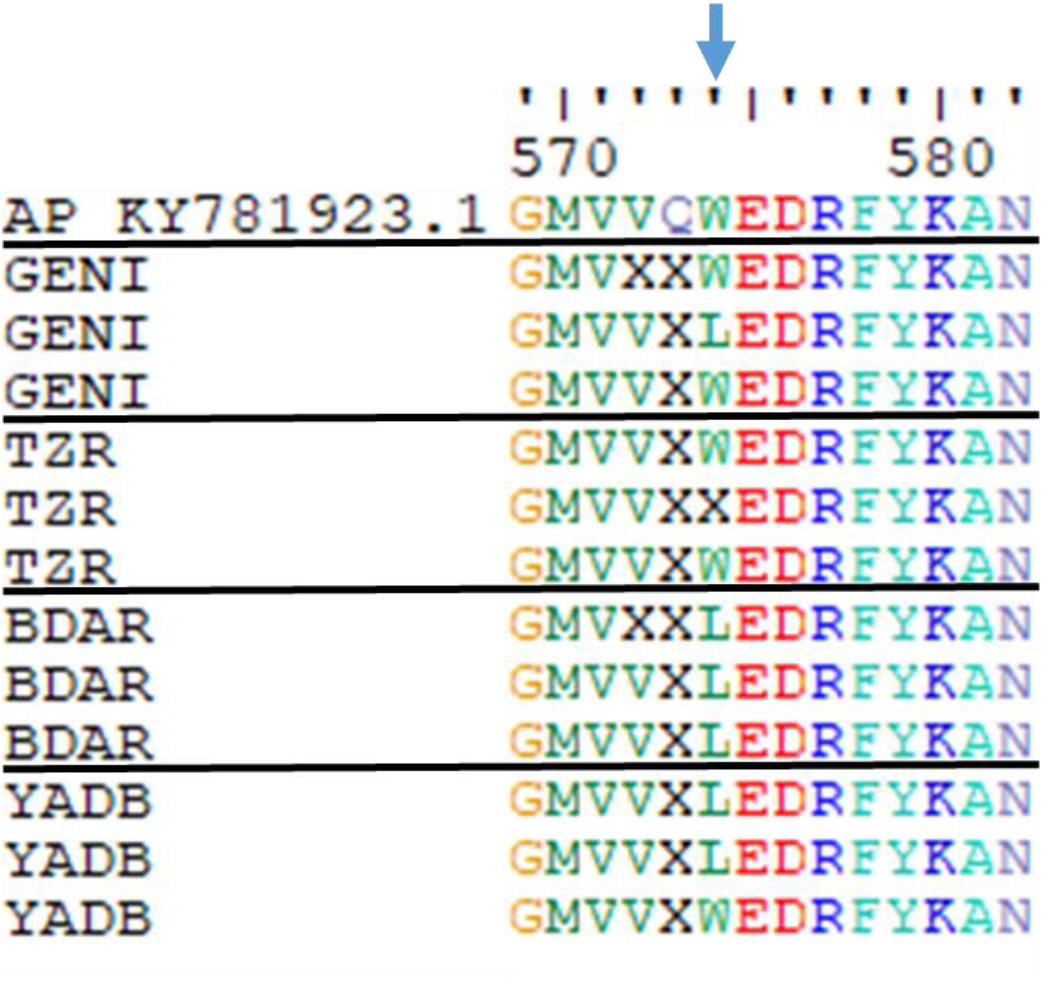
Alignment of partial ALS gene sequence of plants from *A. palmeri* resistant populations. Position 574 is marked with a blue arrow. Positions refer to the known ALS gene sequence of *A. palmeri* (KY781923.1).

## 4 DISCUSSION

*Amaranthus palmeri* was first documented in Israel in the late 1950s, however, its increasing distribution alongside herbicide resistance evolution has marked this species as a major pest in Israeli agriculture in the last 15 years. Recent reports on the occurrence of *A. palmeri* in several other Mediterranean countries (e.g., Spain, Greece, Turkey, and Italy) have raised the concern of the European weed research community. The CLIMEX model, developed by Kistner and Hatfield (2018), predicted the spread of *A. palmeri* in Europe, including most of France, Italy, and south-eastern Europe, and even indicated that the species may spread to northern European countries. According to the updated CLIMEX model recently published by EPPO (2020), the regions most vulnerable to *A. palmeri* invasion under current and future climate scenarios are the Mediterranean and Anatolia. Moreover, the current distribution of *A. palmeri* in the Americas may prove that this species is highly adaptable to various climatic conditions (Roberts & Florentine, 2022). Apart from its native range, *A. palmeri* may be found in the US, Dominican Republic, Brazil, Uruguay, and Canada. Thus, we expect further distribution of this species to more European countries shortly.

In Israel, *A. palmeri* was discovered in various habitats, however, most of the populations were in agricultural fields within crop rows. This may emphasize the opportunist nature of this species, as it benefits from the agricultural practice of providing water and nutrients to aid its growth and expand its distribution using Ag. machinery. As a summer, C4, fast-growing weed in an arid/semi-arid environment, *A. palmeri* has an advantage compared to native weed species and C3 crop plants. In Texas, it was found that *A. palmeri* has an extended emergence period in the field, and even late–emerging individuals may deposit significant amounts of seeds in the soil seedbank (Werner et al., 2020). In addition, fast growth in the early season may enhance competition and thus reduce overall crop plant growth and yield production. Indeed, in Kansas, Horak & Loughin (2000) reported that during June, emerging *A. palmeri* height increased from 0.18 to 0.21 cm growing-degree-day^-1^ and reached maximum heights greater than 2 m. A second cohort emerged approximately three weeks later, with height increases of 0.11 to 0.17 cm growing-degree-day^-1^ and a maximum height of 174 cm.

Apart from its vast distribution, *A. palmeri* showed an extraordinary ability to evolve herbicide resistance to various MOA (Yanniccari et al., 2022). With 67 herbicide resistance reported cases in North America alone, this species jeopardizes agricultural productivity in many countries. Recent reports of herbicide resistance populations from Turkey, Greece, Spain, and Italy support the problematic nature of this species (Kanatas et al., 2021; Manicardi et al., 2023; Mennan et al., 2021; Milani et al., 2021). In our study, we tested the response of plants from *A. palmeri* populations to both glyphosate and trifloxysulfuron. Recent reports from Spain and Turkey have indicated the occurrence of glyphosate-resistant populations of *A. palmeri* in Europe (Manicardi et al., 2023; Mennan et al., 2021). Due to the short period between reports on the first identification of this species in these countries and those of herbicide resistance, one may assume that these populations were glyphosate-resistant upon introduction. However, further research, including population genetics and distribution analysis, is needed to prove this theory.

For all tested populations, glyphosate resistance was not detected, although a high survival rate was recorded following exposure to half of the recommended labeled field rate in several populations (Fig. 1A). Further selection pressure may increase the occurrence of glyphosate-resistant individuals within these populations, and should be taken into account by herbicide application programs in the area. As mentioned above, glyphosate resistance was already documented in Europe, however, we may assume that the different origins of the *A. palmeri* introduced populations may explain the distinct herbicide responses. Moreover, *A. palmeri* was reported in Israel over 60 years ago (Danin, 2022), while in Spain and Turkey, this species was observed only after 2000. This supports the theory of different introduction origins for *A. palmeri* between countries. In addition, the fact that glyphosate-resistant crops are not grown in these European countries, as well as in Israel (Rubin & Matzrafi, 2015), reinforces the assumption of the introduction of glyphosate-resistant populations.

Resistance for the ALS inhibitor trifloxysulfuron was highly abundant among the tested populations (Fig. 2). High resistance levels were recorded for populations originating from the Southern Coastal Plain, with a 0.77 R-value for the region (Table 3). Within the region, five out of seven tested populations at the Southern Coastal Plain showed an R-value of ∼0.9 in response to the full rate of trifloxysulfuron treatment (7.5 g ai ha^-1^; Table 3). One possible explanation for this trend may be that this area is the main region for corn and cotton grown in Israel (Rubin & Matzrafi, 2015). The application of ALS inhibitors such as foramsulfuron, pyrithiobac-sodium, rimsulfuron, and trifloxysulfuron is widespread for both crops. In cotton, trifloxysulfuron and pyrithiobac-sodium may be applied several times in one season, depending on crop growth rate and weed infestation levels. Moreover, the main weeds for both crops are *Amaranthus* species, with an emphasis on *A. palmeri* and *A. tuberculatus* (Rubin & Matzrafi, 2015). Thus, we may assume that a strong selection pressure exerted by these herbicides is the main reason for the high occurrence of *A. palmeri* resistance evolution. Resistance to ALS inhibitors was already reported in Israel and other Mediterranean countries since the early 2000s (Heap, 2023). However, this is the first study documenting the extent of this phenomenon across the country. Our results indicate that the R-value for all regions is higher than 0.5, highlighting the abundance of trifloxysulfuron resistance (Table 3). Moreover, our dose-response studies suggest that some populations show resistance to up to four times the recommended labeled field rate of trifloxysulfuron (30 g ai ha^-1^; Fig. 3).

Previous studies in populations of *A. palmeri* from Spain (Torra et al., 2020) and Italy (Milani et al., 2021), have shown that the substitution of Trp574 to Leu usually correlates with resistance to SU and IMI herbicicdes. In other weed species such as *Raphanus raphanistrum*, this mutation can even confer resistance to all five groups of ALS herbicides (Tan & Medd, 2002). However, for *A. palmeri*, substitution in this position have shown resistance to PTB, IMI, SU and TP but not to SCT inhibitors (Singh et al., 2019). In our study, a substitution in position 574 from Trp to Leu, was found in all populations (Fig. 4) and mutation frequency was higher for populations BDAR and YADB.

Our results may suggest that the use of trifloxysulfuron and other ALS inhibitors to control *A. palmeri* infestation in these fields may be ineffective. In other field crops, such as tomato and peanut, using ALS inhibitors (e.g., rimsulfuron, imazamox, and imazapic) is a common agricultural practice. It is well documented that resistance to one ALS inhibitor may predict failure in response to other herbicides from the same chemical family and even others from the same MOA (Heap, 2023). Thus, the documented resistance to trifloxysulfuron may influence the management of *A. palmeri* in other field crops.

## AUTHOR CONTRIBUTION

JA-N and MM designed the experiments. JA-N, AW, HE and MM have collected the seeds of A. palmeri. JA-N and MM conducted herbicide response assays. JA-N, EW, and MM analyzed the data. JA-N, AW, EW, HE, and MM wrote and reviewed the manuscript. All authors have read and approved the manuscript.

## ACKNOWLEDGEMENTS

The authors would like to thank Shaharit Ziv, Daniel Neta, Neta Solomon, and Sahar Malka for their valuable assistance.

---

**Figure S1.** Representative images of a dose-response experiment showing the phenotypic response of glyphosate-treated *A. palmeri* plants from five populations; KAV-4, YIZ, EVE, 77R, and TZR (A-E). The recommended labeled field rate = 720 g ae ha^-1^, marked in red. n=10. Effect of increasing glyphosate application rates on plants’ shoot fresh weight (F) and survival (G) from five selected *A. palmeri* populations.

**Figure S2.** Effect of increasing application rates of trifloxysulfuron on the shoot fresh weight of five selected *A. palmeri* populations over two experimental runs (GENI, TZR, BDAR, YADB, and GE’A; A and B). The recommended labeled field rate = 7.5 g ai ha^-1^. n=10.

