## Supplementary material for "Occurrence of *Amaranthus palmeri* in Israeli agriculture: status of spread and response to glyphosate and trifloxysulfuron"

**Table S1.** Dose-response parameters for sensitive (KAV-4) and resistant-suspected (EVE, YIZ, 77R, and TZR) *A. palmeri* populations tested for their response to glyphosate.

| | **Pop ID** | **ED$_{50}$ (g ae ha$^{-1}$)** | **SE** | **LD$_{50}$ (g ae ha$^{-1}$)** | **SE** |
|---|---|---|---|---|---|
| **Experimental run No. 1** | | | | | |
| | KAV-4 | 57.2206 | 10.1924 | 189.082 | 0 |
| | EVE | 62.1932 | 6.9337 | 102.555 | 0 |
| | YIZ | 3.9408 | 8.2625 | 165.491 | 54.522 |
| | 77R | 92.3636 | 12.3417 | 189.286 | 124.528 |
| | GENI | 79.4562 | 13.5876 | 151.634 | 21.072 |
| **Experimental run No. 2** | | | | | |
| | KAV-4 | 84.1833 | 6.6971 | 134.26 | 173.894 |
| | EVE | 58.3492 | 6.1131 | 87.699 | 65.654 |
| | YIZ | 62.1697 | 16.7261 | 50.268 | 15.736 |
| | 77R | 58.3492 | 4.6154 | 211.576 | 39.49 |
| | GENI | 63.8026 | 5.9033 | 171.749 | 21.821 |

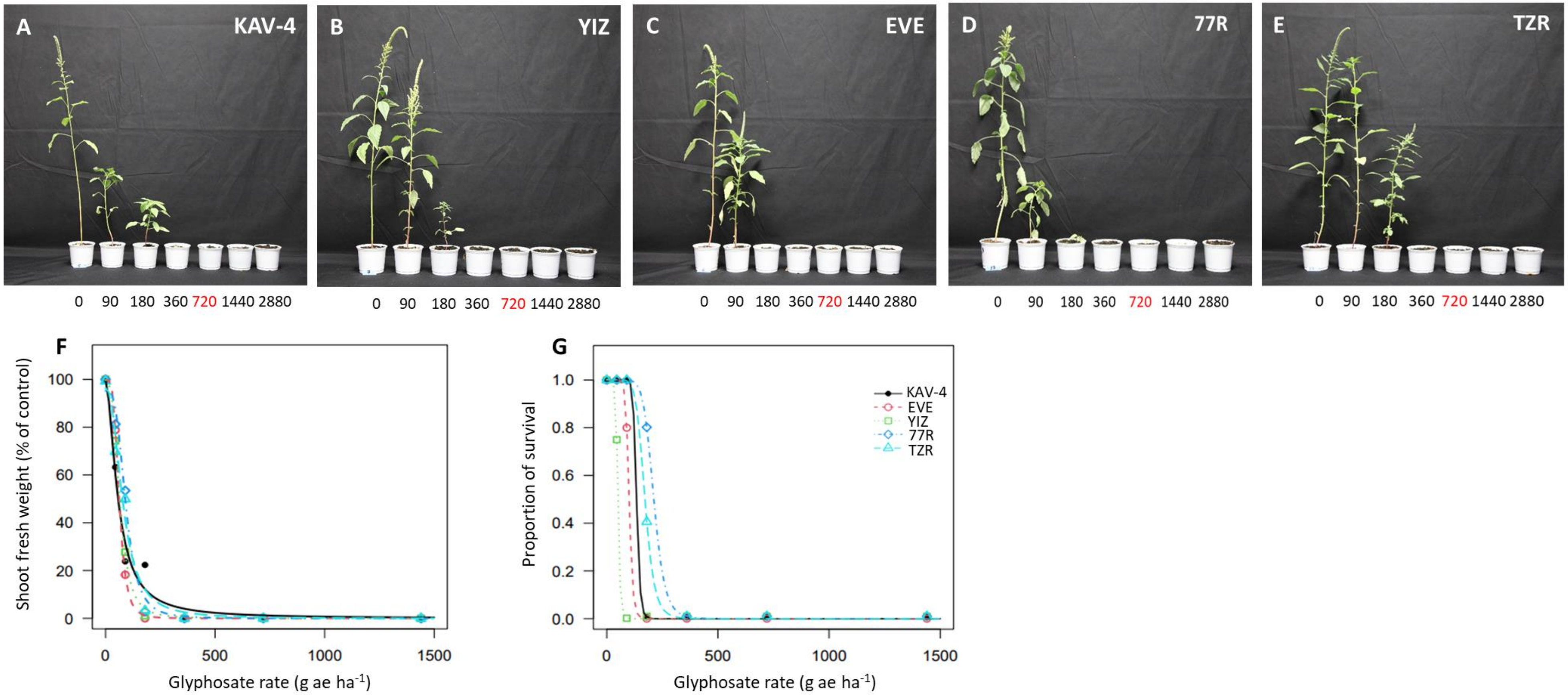
A
KAV-4
B
YIZ
C
EVE
D
77R
E
TZR
0 90 180 360 720 1440 2880
F
Shoot fresh weight (% of control)
Glyphosate rate (g ae ha-1)
G
Proportion of survival
KAV-4
EVE
YIZ
77R
TZR
Glyphosate rate (g ae ha-1)

Fig. S1

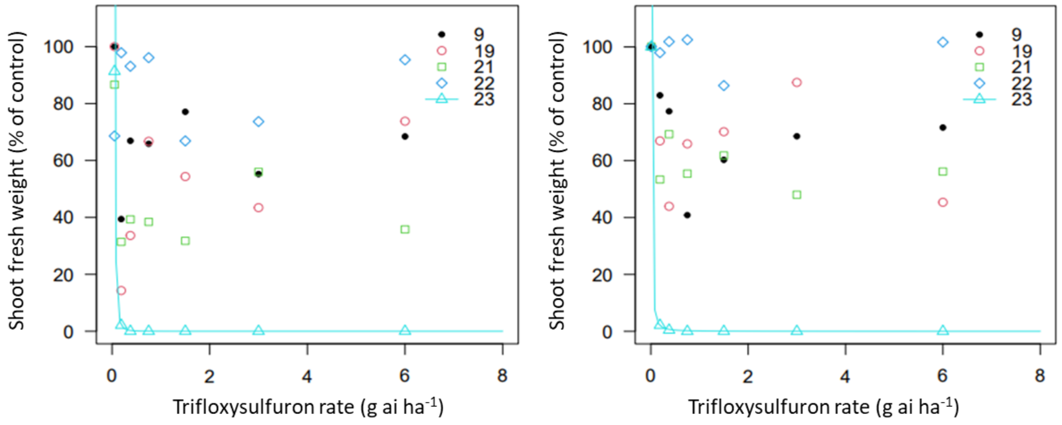
Shoot fresh weight (% of control)
Trifloxysulfuron rate (g ai ha-1)
9
19
21
22
23
Shoot fresh weight (% of control)
Trifloxysulfuron rate (g ai ha-1)
9
19
21
22
23

Fig. S2
